# The evolution of infanticide by females in mammals

**DOI:** 10.1101/405688

**Authors:** Dieter Lukas, Elise Huchard

## Abstract

In most mammalian species, females regularly interact with kin, and it may thus be difficult to understand the evolution of some aggressive and harmful competitive behaviour among females, such as infanticide. Here, we investigate the evolutionary determinants of infanticide by females by combining a quantitative analysis of the taxonomic distribution of infanticide with a qualitative synthesis of the circumstances of infanticidal attacks in published reports. Our results show that female infanticide is widespread across mammals and varies in relation to social organization and life-history, being more frequent where females breed in groups and have intense bouts of high reproductive output. Specifically, female infanticide occurs where the proximity of conspecific offspring directly threatens the killer’s reproductive success by limiting access to critical resources for her dependent progeny, including food, shelters, care or a social position. In contrast, infanticide is not immediately modulated by the degree of kinship among females, and females occasionally sacrifice closely related juveniles. Our findings suggest that the potential direct fitness rewards of gaining access to reproductive resources have a stronger influence on the expression of female aggression than the indirect fitness costs of competing against kin.

## Introduction

Recent work has emphasized that competitive strategies of female mammals are often strikingly symmetrical to those observed in males, including displays and ornaments, fighting and weaponry, dominance hierarchies, and reproductive suppression [Clutton-Brock 2007, 2013; Stockley & Bro-Jørgensen 2011; Clutton-Brock & Huchard 2013]. However, while interactions among conspecific male mammals are often contextual and temporally limited to competition over access to mating partners [Emlen & Oring 1977; Connor & Krützen 2015], interactions among females tend to occur across extended periods and multiple settings [Gowaty 2004; Clutton-Brock 2016]. Females may thus compete over a diversity of resources - including food, resources necessary to breed (burrows, home-range), or offspring care [Clutton-Brock 2007]. In addition, since female mammals are typically philopatric, they may often compete with kin. It has therefore proven difficult to identify the determinants of overt female-female competition, in particular in societies that are structured around female kinship. The challenge has been to understand how the direct benefits of competition may be balanced with the indirect fitness costs of competing against kin, especially in the case of extremely harmful behaviour such as infanticide [Clutton-Brock et al. 2001; Young et al. 2006].

The killing of rivals’ offspring represents one violent manifestation of intrasexual competition, and a significant source of juvenile mortality in some populations [Palombit 2012], with adults of both sexes committing infanticide. It has been intensely studied in male mammals, where fifty years of field research have shown that it has evolved as a sexually selected strategy over access to mating partners [Hrdy 1974]. In cases where the presence of a dependent offspring prevents the mother from becoming pregnant again, committing an infanticide allows the killer to create extra reproductive opportunities. This strategy is particularly common in polygynous societies where one or a few alpha male(s) monopolize mating opportunities over short periods before losing dominance to others [van Schaik & Janson 2000; Lukas & Huchard 2014; Palombit 2015]. Since in most instances infanticide is committed by males who recently joined the group, they are unlikely to kill any related offspring [Hrdy 1979]. In contrast, as for other forms of female competition, little is known about the determinants and consequences of infanticide by females other than the mother, although it may be more prevalent than infanticide by males, both within and across taxa [Hrdy 1976, Blumstein 2000; Digby 2000]. Unlike males, female killers do not benefit from extra mating opportunities [Agrell et al. 1998; Digby 2000], because male mammals generally do not invest into offspring care to the extent that it would prevent them from mating with other females [Kleiman & Malcom 1981]. If anything, killing a dependent juvenile may exacerbate female mating competition by speeding-up the resumption to fertility for the mother of the victim. Several plausible scenarios explaining the occurrence of infanticide by females have been proposed based on a synthesis of natural observations [reviewed by Digby 2000]. Symmetrically to the patterns observed for male infanticide, predation for nutritional gains (H1: ‘exploitation’ hypothesis) may not provide a general explanation for female infanticide as killers have relatively rarely been observed to consume victims partially or entirely (e.g. [Goodall 1986; Blumstein 2000]). Instead, hypotheses regarding the adaptive benefits of female infanticide have focused on how killings might facilitate access to resources that are critical to successful reproduction (H2: ‘resource competition’ hypotheses) [Digby 2000]. Female killers might be defending access to an exclusive territory or shelter, or attempting to expand their breeding space when they target victims outside their home-range (H2.1: ‘breeding space’ hypothesis) (as in black-tailed prairie dogs [Hoogland 1985] or Belding’s ground squirrels [Sherman 1981]). In species where females only associate temporally to breed, killers may defend access to their own milk, by discouraging attempts to suckle from unrelated juveniles (H2.2: ‘milk competition hypothesis) (as in Northern elephant seals: [Le Boeuf et al. 1973]). In species who breed cooperatively, killers may defend access to extra offspring care by group mates other than the mother by altering the helper-to-pup ratio in their own group (H2.3: ‘allocare’ hypothesis) (as in meerkats [Clutton-Brock et al. 2001; Young et al. 2006], banded mongooses [Gilchrist 2006; Cant et al. 2014], or marmoset [Digby 1995]). Finally, in species who live in stable groups, killers may defend their offspring’s future social status (in species with stable hierarchies) or group membership (in species with forcible evictions) by eliminating future rivals (H2.4: ‘social status’ hypothesis) (as in some Old World primates [Hrdy 1976; Digby 2000]). Finally, it is unclear whether females might refrain from committing infanticide when this is likely to lead to indirect fitness costs. If so, they might be less likely to commit infanticide in species where most interacting females are related, or they might be able to target unrelated offspring exclusively. Alternatively, the benefits of competition in any or all these circumstances might be high enough to outweigh any potential indirect fitness costs.

Here, we present an investigation of the distribution and circumstances of infanticide by female mammals, based on data from 289 species from across 14 different Orders collected from the primary literature. The combination of a quantitative synthesis of the taxonomic distribution of infanticide with a qualitative analysis of the circumstances of infanticidal attacks (including traits of the killer and victim) can contribute to reveal the ecological, life-history or social determinants of female reproductive competition across mammalian societies, and their relevance to the occurrence of female associations and interactions within and among matrilines. Our aim is to provide a starting point for the investigation of the likely causes and situations under which female infanticide occurs, bearing in mind that our analytical framework suffers from several caveats. First, infanticide is uncommon and difficult to observe, so that some species might wrongly be classified as ‘non-infanticidal’. Second, the analyses rely on a rough categorization of the circumstances of infanticide and may oversimplify its determinants, which could be influenced by multiple factors in a given species. We first summarize the social organisation and life-histories of species in which infanticide by females has been observed, in order to evaluate the conditions under which infanticide is most common (Table 1). In particular, we investigate whether philopatry and average relatedness among groups of interacting females reduce the occurrence of female infanticide. Next, we perform specific phylogenetic analyses to test core predictions generated by each hypothesis and assess whether females have been observed to commit infanticide in species in which they are most likely to benefit from such killings. In addition, we investigate whether population-level information on the traits of killers and victims are compatible with predictions generated by each hypothesis. All our predictions and tests are summarized in Table 2. We first show that, across all species, the distribution and occurrence of infanticide by females is better explained by resource competition than by exploitation. We next test support for each for the four resource-competition hypotheses. We assess whether (i) instances where females kill offspring in neighbouring ranges (‘breeding space hypothesis’) are most likely explained by competition over breeding space; whether (ii) instances where females kill offspring born in the same breeding association by competition over milk (‘milk competition hypothesis’); whether (iii) instances where females kill offspring in groups where usually only a single female reproduces by competition over offspring care (‘allocare hypothesis’); and whether (iv) instances where females kill offspring born in groups with multiple breeders by competition over social status or group membership (‘social status hypothesis’).

**Table 1:**
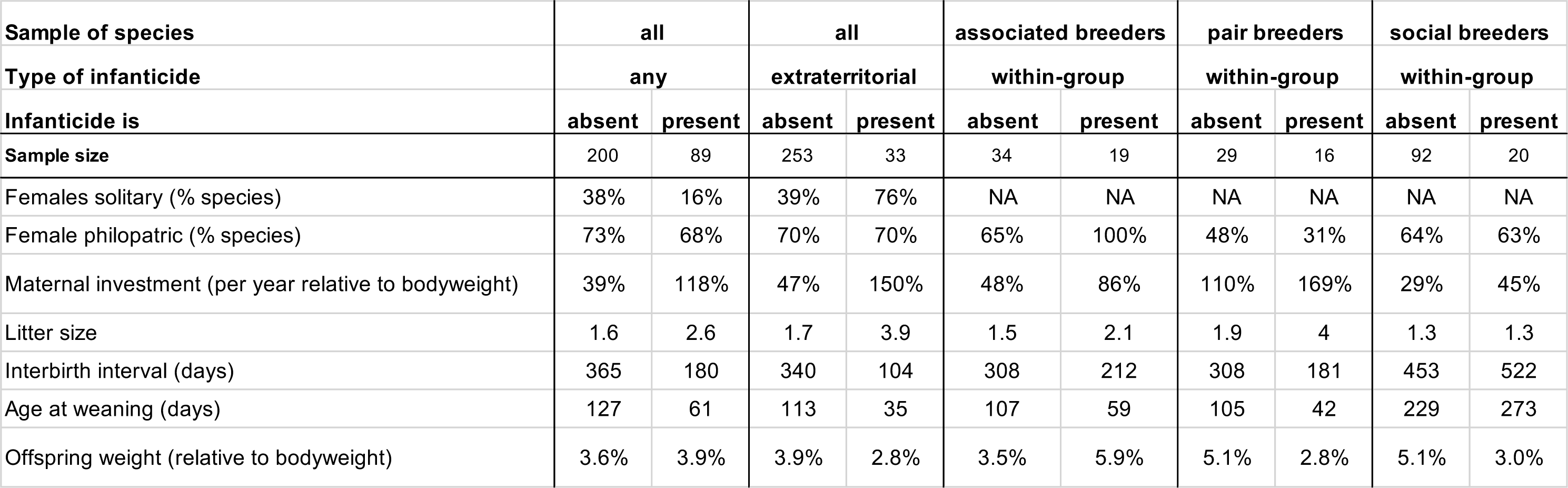
The social and life-history conditions associated with the distribution of infanticide by females. Females are more likely to commit infanticide in group-living species, but whether breeding females are philopatric or disperse does not appear to be associated with the distribution of infanticide. In general, species with infanticide by females are characterized by high maternal investment, which is either expressed as females having large litters, short lactational and interbirth periods, and/or relatively large offspring.

| **Sample of species** | **all** |  | **all** |  | **associated breeders** |  | **pair breeders** |  | **social breeders** |  |
| --- | --- | --- | --- | --- | --- | --- | --- | --- | --- | --- |
| **Type of infanticide** | **any** |  | **extraterritorial** |  | **within-group** |  | **within-group** |  | **within-group** |  |
| **Infanticide is** | **absent** | **present** | **absent** | **present** | **absent** | **present** | **absent** | **present** | **absent** | **present** |
| **Sample size** | 200 | 89 | 253 | 33 | 34 | 19 | 29 | 16 | 92 | 20 |
| Females solitary (% species) | 38% | 16% | 39% | 76% | NA | NA | NA | NA | NA | NA |
| Female philopatric (% species) | 73% | 68% | 70% | 70% | 65% | 100% | 48% | 31% | 64% | 63% |
| Maternal investment (per year relative to bodyweight) | 39% | 118% | 47% | 150% | 48% | 86% | 110% | 169% | 29% | 45% |
| Litter size | 1.6 | 2.6 | 1.7 | 3.9 | 1.5 | 2.1 | 1.9 | 4 | 1.3 | 1.3 |
| Interbirth interval (days) | 365 | 180 | 340 | 104 | 308 | 212 | 308 | 181 | 453 | 522 |
| Age at weaning (days) | 127 | 61 | 113 | 35 | 107 | 59 | 105 | 42 | 229 | 273 |
| Offspring weight (relative to bodyweight) | 3.6% | 3.9% | 3.9% | 2.8% | 3.5% | 5.9% | 5.1% | 2.8% | 5.1% | 3.0% |

**Table 2:** Testing the core predictions generated by the different hypotheses proposed to explain the distribution of infanticide by females. For each of the main hypotheses, we tested two core predictions in phylogenetic comparisons and two predictions about the individual traits from the field observations. For the comparisons, we list the sample of species included.

| Hypothesis | Type of infanticide /<br>Sample of species | Core prediction(s) | Support? | Predictions of individuals'<br>characteristics | Support? |
| --- | --- | --- | --- | --- | --- |
| **H1: exploitation** | *any form of infanticide* | - primarily in carnivores | No | - killer: any reproductive state | No |
|  | *across all species* | - infanticide also by males | No | - victim: any age | No |
| **H2: resource competition** | *any form of infanticide* | - harsher environments | Yes | - killer: gestating/lactating | Yes |
|  | *across all species* | - higher maternal investment | Yes | - victim: dependent on care | Yes |
| **H2.1: over breeding space** | *extraterritorial infanticide* | - more burrow use | Yes | - killer: gestating/lactating | Yes |
|  | *across all species* | - exclusive home-ranges | Yes | - victim: unweaned | Yes |
| **H2.2: over milk** | *within group infanticide* | - higher energetic milk content | No | - killer: lactating | Yes |
|  | *across associated breeders* | - faster offspring growth | No | - victim: attempting to nurse | Yes |
| **H2.3: over allomaternal care** | *within group infanticide* | - allocarers present | Yes | - killer: gestating/lactating | Yes |
|  | *across pair breeders* | - more helpers/competitors present | Yes | - victim: dependent on care | Yes |
| **H2.4: over social status** | *within group infanticide* | - nepotistic hierarchy present | Yes | - killer: high social rank | Yes |
|  | *across social breeders* | - evictions occur | Yes | - victim: any age | Yes |

## Materials & Methods

Following Digby [2000], we use a broad definition of infanticide as an act by one or more non-parents that makes a direct or significant contribution to the immediate or imminent death of conspecific young. This definition excludes instances where mothers kill their own offspring (which are considered to result from parent-offspring conflict [O’Connor 1978] rather than from intrasexual competition) and includes cases where infants die as the result of the physical aggression (direct infanticide) as well as cases where the enforced neglect of an infant, such as kidnapping, ultimately causes death (indirect infanticide). Although the latter cases are often excluded from studies of infanticide due to their proximate form of ‘overzealous’ allomaternal care [Hrdy 1976], their ultimate consequence - infant death - contributes to shape their evolution as infanticidal behaviour. We included infanticide records from both wild and captive populations for which the killer was unambiguously identified as an adult female. For each species, we recorded whether observations occurred in a captive setting or under natural conditions. Species for which no case of infanticide has ever been observed were included only if detailed observations on individual females and juveniles were available, either from repeated captive observations or from field studies occurring across at least three reproductive seasons, to minimize the risk of misclassifying them as “non-infanticidal”. Given that infanticide is difficult to observe, we focused on studies that were performed in circumstances under which it could be expressed and detected (co-housing of multiple females; detailed observations or specific females from before birth until weaning) or on studies that followed the fate of one or more offspring cohort(s) and recorded the causes of juvenile mortality. Data were collected from searches through the scientific literature, starting with major reviews on the topic of female infanticide [Wasser & Barash 1983; Agrell et al. 1998; Ebensperger 1998; Blumstein 2000; Digby 2000; Ebensperger & Blumstein 2007; Stockley & Bro-Jorgensen 2011; Clutton-Brock & Huchard 2013] and performing backward and forward citation searches to identify relevant observations. While we might not have included all species in which the status of infanticide by females is documented (i.e., known to be either absent or present), we believe that this search strategy should not lead to systematic biases with regards to the tested hypotheses. We repeated all analyses excluding the 42 species for which we only found observations in captive settings (27 species with and 15 species without female infanticide), which did not change any of the results.

For the comparative analyses, we extracted data for each species on variables linked to the four different hypotheses (see Table 2). From published databases, we obtained information on: social organisation (classified as: solitary breeders [breeding females have exclusive ranges in which they do not tolerate any other breeding individual], pair breeders [home-ranges contain a single breeding female and a single breeding male but may contain additional non-breeding individuals], associated breeders [females share the same space for breeding but associations are unstable and tend not to last beyond the breeding season], or social breeders [several breeding females share the same home-range across multiple breeding seasons]) [Lukas & Clutton-Brock 2017]; female philopatry and dispersal (whether most breeding females have been born in their current locality/group or elsewhere) [Lukas & Clutton-Brock 2011]; carnivory (whether the diet of a species includes meat or not) [Wilman et al. 2014]; infanticide by males (whether males have been observed to kill conspecific young) [Lukas & Huchard 2014]; environmental climatic harshness (a principal component, with high values indicating that rainfall is low and temperatures are cold and unpredictable across the known range of a species) [Botero et al. 2014]; maternal investment (mean body size of offspring at weaning multiplied by the mean number of offspring per year, divided by mean body mass of adult females) [Sibly et al. 2014]; the use of burrows or nest holes for breeding (information was taken from the papers used to extract information on the absence or presence of infanticide by females); litter size (number of offspring per birth); offspring mass at birth (grams); weaning age (age at which offspring are independent, in days); inter-birth interval (time between consecutive births, in days) [Jones et al. 2009]; energetic value of milk (MJ/ml based on the protein, sugar, and fat composition) [Langer 2008; Barton & Capellini 2011; Hinder & Milligan 2011]; offspring care by fathers and/or non-parental group members (whether offspring receive milk or food from or are being regularly carried by group members who are not the mother) [Lukas & Clutton-Brock 2017]. In addition, we completed information obtained from these databases by collecting extra data from the primary literature on dominance hierarchies and mechanisms of rank acquisition in social groups (whether all adult females can be arranged in a dominance hierarchy and if so, whether an individual’s rank is influenced by age and/or nepotism); and forcible evictions (whether females use aggression to exclude other females from their own social group). For each species in which females had been observed to kill conspecific young, we used the primary literature to record as much information as possible regarding the characteristics of the killer (age and reproductive state) and of the victim (age, sex, and relatedness to killer) to test specific predictions. The full dataset is provided in Supplementary Table 1 (comparative data) and Supplementary Table 2 (individual characteristics data), with all references for data specifically collected here in Supplementary File 1.

In addition to performing comparisons assessing contrasts in the presence or absence of infanticide across all species in our sample, we classified species into different types according to each of the four resource competition hypotheses (Table 2). This classification also aimed at controlling for a potential confounding effect of social organization, as our analysis might otherwise detect factors associated with the evolution of sociality if females are more likely to have been observed to commit infanticide in some social systems. For the breeding space hypothesis, we only included instances of infanticide in which females did not share a home-range with the mother of the victims; for the milk competition hypothesis, we restricted the sample to associated breeders; for the allocare hypothesis, we only included pair; and for the social competition hypothesis, we only looked at social.

For the comparative analyses, the phylogenetic relatedness between species was inferred from the updated mammalian supertree [Rolland et al. 2014]. We fitted separate phylogenetic models using MCMCglmm [Hadfield & Nakagawa 2010] to identify the extent to which each of the predicted variables (Table 2) explains the presence of infanticide by females across species (binary response, assuming a categorical family of trait distribution). Following the recommendations of Hadfield [2010], we set the priors using an uninformative distribution (with variance, V, set to 0.5 and belief parameter, nu, set to 0.002). Each model was run three times for 100,000 iterations with a burn-in of 20,000, visually checked for convergence and for agreement between separate runs.

## Results

### Social organisation and infanticide by females

Infanticide by females has been observed in 89 (31%) of the 289 mammalian species in our sample (Table 1). Female infanticide (of any type) varies with the social organisation and is more frequent when females breed in groups (Figure 1): it has been observed in 43% of associated breeders, in 36% of pair breeders, and in 30% of social breeders, but only in 18% of solitary breeders. Across all species, females are equally likely to kill offspring when they are philopatric (47 of 135 species, 34%) than when they disperse to breed (17 of 59 species, 29%) (effect of female dispersal on presence of infanticide by females: −10.1, 95% CI −39.3 – 11.3, p=0.34) but there are differences for two types of social organisation: across associated breeders infanticide only occurs in philopatric species; while across pair breeders infanticide is more likely to occur in species in which females disperse (Table 1). Across all group-living species (associated breeders, pair breeders with helpers, social breeders), there is no relationship between levels of average relatedness among female group members and whether infanticide by females of offspring born in the same group does (median levels of average relatedness across 10 species 0.09, range 0.01-0.38) or does not occur (median levels of average relatedness across 24 species 0.21, range −0.03-0.52) (effect of levels of average relatedness on presence of infanticide by females: −21.1, 95% CI −81.4 – 10.1, p=0.18). Across species in which groups are stable (i.e., excluding associated breeders where groups are sometimes difficult to define and can be very large), levels of average relatedness are slightly higher when infanticide occurs (see also [Lukas & Clutton-Brock 2018]). The population-level information show instances of killers being close kin of the victim in 33% of species (22/65), with either grandmothers killing their grandchildren or aunts killing their nieces, with kin being common victims in cooperative breeders (8 of 12 cooperative breeders) but also in several social breeders (14 of 51 social breeders).

**Figure 1:**
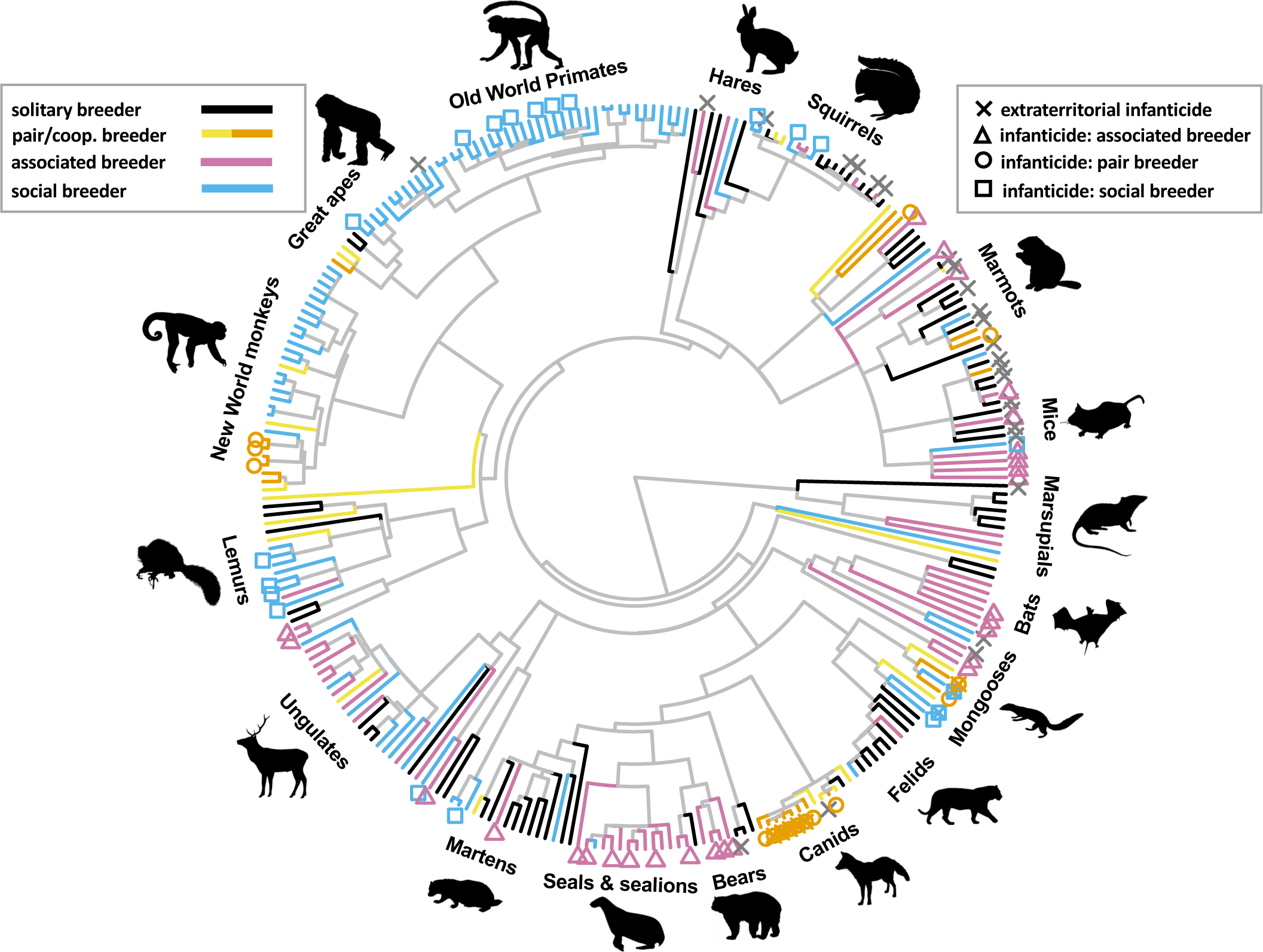
The distribution of the different forms of infanticide by females in relation to the social organisation across The mammalian species in our sample. (for picture credit see Supplementary File 2).

### Life-histories and infanticide by females

Energetic investment into reproduction by mothers is higher in species with any form of infanticide by females compared to the remaining species (Table 1). However, these patterns encountered across all species do not reflect a general association between infanticide and all these traits, but rather the fact that each different scenario of infanticide is associated with specific life histories, with species having either larger offspring mass at birth, shorter time to weaning and between births, and/or larger litter sizes. Females in infanticidal species where there is usually a single breeding female per home range (extraterritorial infanticide and infanticide in pair breeders) have larger litters than females in species without infanticide (Table 1). Species in which females kill offspring in a breeding association are characterized by fast-growing offspring, while offspring are relatively small at birth in species in which females kill offspring in stable groups (Table 1).

#### H1: Exploitation

We find no support for predictions suggesting that females kill conspecific offspring primarily for exploitation (Table 2). Across species, infanticide by females is as likely to occur in the absence of infanticide by males (44 of 147 species, 30%) as in its presence (43 of 135 species, 32%) (effect of presence of male infanticide on presence of female infanticide 2.1, 95% CI −13.5 – 16.9, p=0.74). Similarly, carnivorous species are not more likely to show infanticide by females (18 of 56 species, 32%) than species in which meat does not constitute an important part of the diet (59 of 192 species, 31%) (effect of carnivory on presence of female infanticide −0.7, 95% CI −11.2 – 10.0, p=0.89). The age of victims varies from birth to beyond independence across species, but is more homogeneous within each type of infanticide (see below), so that killings do not appear simply opportunistic.

#### H2: *Resource competition*

Infanticide by females appears more likely to occur where competition over resources is expected to be more intense (Table 2). The climatic environments of species in which females commit infanticide are harsher (as estimated by a principal component reflecting low unpredictable rainfall and cold seasons) than the environments of species in which infanticide has not been observed (effect of environmental harshness on presence of female infanticide 7.0, 95% CI −0.2 – 14.7, p=0.03, 54 species with infanticide and 193 without) (Figure 2a). In species where females commit infanticide, they invest substantially more energy into the production of offspring, being able to produce the equivalent of 1.0 times their own body mass in offspring mass per year (number of offspring times mass of weaned offspring; median across 41 species, range 0.05 – 12.1 times) compared to 0.33 times in species in which infanticide has not been observed (median across 77 species, range 0.03 – 11.1) (effect of maternal energetic investment on presence of female infanticide 25.8, 95% CI 1.5 – 58.7, p<0.001) (Figure 2b).

**Figure 2:**
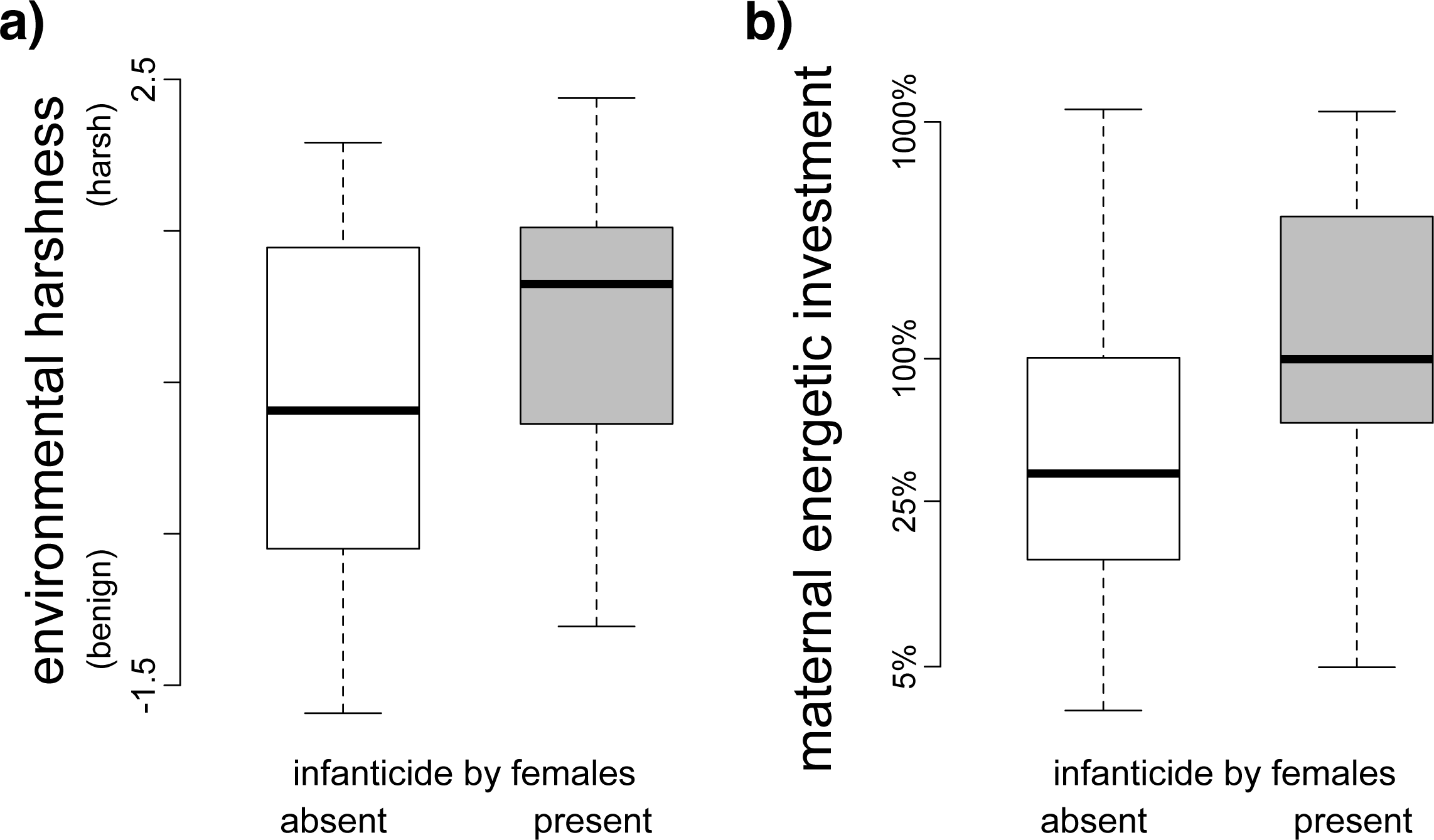
Factors associated with female competition and the distribution of infanticide by females. Species in which female infanticide is present are, on average, characterized by (a) living in harsher environments (lower and more unpredictable rainfall and temperatures) and by (b) higher maternal energetic investment (total mass of weaned offspring produced per year relative to maternal mass). Black lines indicate the median across the species in the sample, boxes contain 75% of the values, and whiskers extend to the extremes

#### H2.1: Competition over breeding space

Thirty-two of the 33 species in which females kill juveniles outside their own home-range keep their offspring in burrows or holes, compared to 93 of the 163 species in which infanticide by females appears absent (effect of burrow use on the presence of infanticide by females 14.4, 95% CI 6.0 – 22.9, p<0.001). The exception is *Semnopithecus entellus*, where “females occasionally steal infants from a neighboring troop” [Hrdy 1976; Mohnot 1980]. In most species in which females kill offspring in neighbouring home-ranges (25 of 33), females generally appear not to tolerate other breeding females close by and most home-ranges only contain a single breeding female (solitary or cooperatively breeding species), while in most other species females form associations or groups (home-ranges contain a single breeding female in 105 out of 268 species) (effect of presence of a single breeding female per home-range on presence of infanticide by females 12.4, 95% CI 0.3 – 30.1, p=0.007). In all cases, the killer was either pregnant or had dependent young of her own (17 species with observations), and all offspring that were killed were not yet weaned (17 species).

#### H2.2: Competition over milk

Across associated breeders, the energy content of milk produced by mothers in species in which females have been observed to kill offspring within the same breeding space (2.8 MJ/100ml, median across 8 species, range 1.2-4.7) does not differ from that of species in which such killings have not been observed (2.1 MJ/100ml, median across 7 species, range 0.8-5.7) (mean effect of milk energy on presence of infanticide by females 2.0, 95% CI −66.3 – 83.1, p=0.97). In associated breeders with female infanticide, offspring do not seem to have greater growth rates (they gain on average 0.28% of their adult body mass per day until weaning, median across 9 species, range 0.04% - 1.72%) than in species in which females have not been observed to kill juveniles (offspring gain on average 0.17% of their adult body mass per day until weaning, median across 11 species, range 0.05% - 0.73%) (effect of presence of infanticide by females on offspring growth rate 6.1, 95% CI −16.2 – 35.4, p=0.59). Killers are either pregnant or have dependent young (21 out of 21 species) and are old, with most reports suggesting that they had been observed to give birth in previous years (they are not primiparous). All victims were reported to be unweaned (22 out of 22 species). In 7 out of 14 infanticide reports from associated breeders, victims were killed as they attempted to suckle from the killer.

#### H2.3 Competition over allomaternal care

Infanticide by females in pair breeders occurs only when fathers provide care (all 16 species) while fathers care for offspring in only 16 of the 39 pair breeding species in which this form of infanticide is absent. In 15 of the 16 pair breeders with female infanticide, additional helpers are present (cooperative breeders), while there are only a further 7 cooperatively breeding species in which females have not been observed to kill juveniles from their own group (female infanticide occurs in 68% [15/22] of cooperative breeders versus in 4% [1/23] of pair breeders in which there are no other helpers). Across species in which offspring receive allocare, the number of potential allocarers is higher in species with, compared to species without female infanticide (infanticide present: 3 allocarers per group, median across 15 species, range 2-23; infanticide absent: 2 allocarers per group, median across 13 species, range 1-20; effect of number of allocarers on presence of infanticide 25.3, 95% CI 1.9 – 48.6, p=0.02). The killer was usually the dominant breeder (as was regularly the case in 9 of 12 species) and was pregnant or with dependent infants in all cases. In most instances the killer and the victim belonged to the same group (11 of 14 species), and were consequently related (10 out of 13 species). Victims were often a few days old and all dependent.

#### H2.4: Competition over social status

Of the 27 social breeders with female philopatry (and available data), female group members do not form social hierarchies in three species, hierarchical rank is determined by age in five species, and rank is influenced by nepotism in 19 species. Females have been observed to kill offspring born to other group members in eight of these latter 19 species, but in none of the species where nepotism does not influence female rank (effect of presence of nepotistic rank acquisition on presence of infanticide by females 134.1, 95% CI 28.8 – 238.1, p<0.001). Females are more likely to kill offspring born to other females in social breeders in which they also aggressively evict other females from their group (infanticide has been observed in 6 of 10 species with evictions and 7 of 35 without evictions) (effect of occurrence of evictions on presence of infanticide by females 7.8, 95% CI −0.2 – 14.4, p=0.02). In all twelve social breeders in which infanticide events have been observed, killers were old and high-ranking. Killers were never pregnant, but in all cases had dependent young of their own. Victims were not yet weaned, and victims might be related to the killer in 5 out of 12 species. There is only one species where the data suggest that females might preferentially kill offspring of one sex: in *Macaca radiata* (a species with female philopatry), female offspring appear to be the predominant victims.

## Discussion

Our findings establish that female infanticide is widespread across mammals and that it is frequently expressed as intensely as competition among males. Females have been observed to kill conspecific juveniles in various species and our comparative analyses provide support to the idea that this behaviour may be adaptive under a wide range of circumstances. Infanticide is more likely to occur in species in which multiple adult females live or breed together than where females breed solitarily, and infanticide appears most frequent in species where females only associate temporarily to breed. Since infanticidal behaviour is relatively rare and may be difficult to detect, this association may reflect the fact that opportunities to commit and to observe infanticides may be greater where females live or breed together. Within each type of social organisation, we do however find that females, like males, appear to commit infanticide when the presence of the victim might otherwise limit their own reproductive success. While infanticide by males has evolved primarily in response to mate competition across mammals [Lukas & Huchard 2014; Palombit 2015], the evolutionary determinants of infanticide by females are apparently more complex, as females may compete over multiple resources.

Several lines of evidence indicate that female infanticide is adaptive, with females killing conspecific offspring in response to competition over resources that are critical for successful reproduction. First, infanticide appears associated with variation in ecology and life-history. Specifically, it is most frequently observed in species facing harsh climatic conditions and making the greatest reproductive efforts; it is unlikely that such associations are due to variations in opportunities to observe or commit infanticides across species. Rather, the potential costs of sharing critical resources might outweigh the risks associated with committing infanticide in such circumstances.

Second, specific determinants of female infanticide identified at the population level by field studies also seem to predict its distribution across species. Extraterritorial infanticides were found to be most frequent in solitary species where females use burrows to give birth and territories to raise offspring, allowing killers to free-up reproductive space for their own offspring. Anecdotal reports suggest that mothers of victims in these species frequently move away after the loss of their offspring [Sherman 1981]. The strong association with burrow use might occur because burrows represent a clear defendable resource, because offspring kept in burrows tend to be altricial [Kappeler 1991] and unable to flee or defend themselves, or because infanticide is more easily observed if researchers know where offspring are. Our findings further show that female infanticide occurs in pair breeders where helpers – fathers or additional group mates – are present. A lower number of helpers per offspring reduces their weight and their chances to survive at independence, such that females might even kill offspring born to close kin, such as their grandchildren [Clutton-Brock et al. 2001; Hodge et al. 2008]. Finally, patterns are slightly more complex in social breeders. There, infanticide preferentially occurs in species where aggressive competition among females leads to the eviction of some individuals – generally young adults - from the group, especially at times when group size increases (e.g. [Kappeler & Fichtel 2012]). In such cases, killing unrelated juveniles may limit future competition and the related risk of being evicted for the killer’s offspring. In addition, in social breeders where females are philopatric, infanticide was only found to occur where female rank acquisition is nepotistic, a hierarchical system where each additional offspring may contribute to strengthen the social status of a matriline – and where infanticide may consequently weaken competing matrilines on the long term.

Anecdotal reports of female pinnipeds killing orphans as they attempted to suckle from them inspired the hypothesis that females compete over milk in species where they only associate to breed [Digby 2000]. While our comparative analyses did not reveal any difference in the energy content of milk of associated breeders in which infanticide is present versus absent, associated breeders nevertheless comprise the species with the highest energy content of milk and the fastest growth rates, and we further found that offspring are larger at birth and weaned at an earlier age in associated breeders with infanticide compared to those without it. The lack of support for the milk competition hypothesis in our analyses may be explained by a noisy dataset, where the absence of infanticide in some species may be due to the fact that it goes undetected if it is hard to observe, or to the evolution of counter-adaptations that protect offspring against infanticide. Alternatively, milk is not the only resource over which these females compete. For example, in the large breeding colonies of pinnipeds, space is sometimes very restricted [Baldi et al. 1996], especially in the immediate vicinity of the harem leaders. These bulls often protect their females and calves from attacks by younger males, and may represent another source of competition for lactating females.

It is likely that, in any given species, infanticide may be triggered by more than one determinant - including some that may not be considered here. A killer may accordingly get multiple benefits from one infanticide event, but may also commit infanticides in more than one context. For example, half of the species of pair breeders committing intra-group infanticides also commit extraterritorial infanticides. It is therefore possible that different types of female infanticide – following our classification - have followed a common evolutionary path. Specifically, it is possible that infanticidal behaviour initially emerged in response to one particular pressure (e.g., competition over access to allocare) in a given species, which subsequently started to express it in other competitive contexts (e.g., competition over breeding territories). However, the limited number of species for which observational data on infanticide are available, as well as heterogeneities in the sample – such as an over-representation of group-living species – introduce uncertainty when attempting to reconstruct the evolutionary history of the trait. It is consequently hard to infer the ancestral state, whether each infanticide type has evolved independently, or how many times infanticidal behaviour has emerged across mammals. Similar difficulties limit causal inferences regarding the association between infanticide and social organization. In some lineages the risk of infanticide might prevent breeding females from forming groups, while in others the evolution of infanticide might be favoured by the constraints or opportunities of a specific social system, in line with patterns observed in males [Lukas & Huchard, 2014].

In addition to the nature of the resources that may directly limit female reproductive success in various types of social organisations [Cavigelli et al. 2003, Saltzman et al. 2009, Rosvall 2011], contrasts in the occurrence of infanticide across species reveal other broad patterns on female reproductive competition in mammals. In particular, the lack of association between female infanticide and philopatry across species (Table 1), as well as a synthesis of observations revealing that killers and victims are commonly related in some contexts, such as in pair breeders where reproductive suppression is common [Lukas & Clutton-Brock 2018], suggest that matrilineality and subsequent increases in average kinship among associated females does not necessarily lead to a reduction in competition among females. Some previous work suggested that mammalian females might be predisposed to behave positively and cooperatively with kin [di Fiore & Rendall 1994], such that species with female philopatry would be characterized by stable social bonds [Silk 2007]. However, the factors leading to limited dispersal and the spatial association of kin frequently also result in high local competition [Frank 1998] which can overcome the potential benefits of cooperation among kin [West et al. 2002]. Studies of competition among males in such circumstances have shown that contrasts in levels of aggression can be explained by variation in the potential direct fitness benefits of winning [West et al. 2001], and it is likely that this also applies to the observed pattern of infanticide by females – where the direct benefits of infanticide in terms of increased access to a critical resource might outweigh its costs, including the indirect fitness costs associated with killing related juveniles.

Our study compiles five decades of behavioural data across species and within populations to elucidate the determinants of infanticide by mammalian females, which are less well understood than those of male infanticide. Our analyses suggest that the distribution of female infanticide across species reflects contrasts in social organisation; infanticide is most frequent in species that breed in groups, which probably have more opportunities for killings and also face greater breeding competition. Female infanticide occurs where the proximity of conspecific offspring directly threatens the killer’s reproductive success by limiting access to critical resources for her dependent progeny, including food, shelters, care or a social position. Finally, these data support the idea that female killers occasionally sacrifice related juvenile conspecifics, and may therefore actively harm their indirect fitness in order to maximize their direct fitness.

## Acknowledgements

We thank Tim Clutton-Brock, Peter Kappeler, Oliver Höner and Eve Davidian for useful discussions.

## Data Accessibility

All data are provided in the supplementary material and are deposited at the Knowledge Network for Biocomplexity doi:10.5063/F1ZG6QFR.

## Authors’ Contributions

Both authors contributed to the conception and design of the study, collected and interpreted the data, wrote the article, and gave final approval; the analyses were done by DL with input from EH.

## Supplementary Material

This submission includes all data in Supplementary Tables 1 (comparative data) and 2 (population observations), with references for observations on infanticide by females in Supplementary File 1, and credits for the animal drawings used in Figure 1 in Supplementary File 2.

## Supplementary File 1 for Lukas & Huchard: The evolution of infanticide by females in mammals

Parts A (citations for comparative data), B (citations for population data), and C (reference list).

### A)Citations for new data in comparative analyses

The table lists the references with the observations on the presence or absence of infanticide by females for each species, and the references with details on the interactions among females in group-living species. All references are listed in full below the second table.

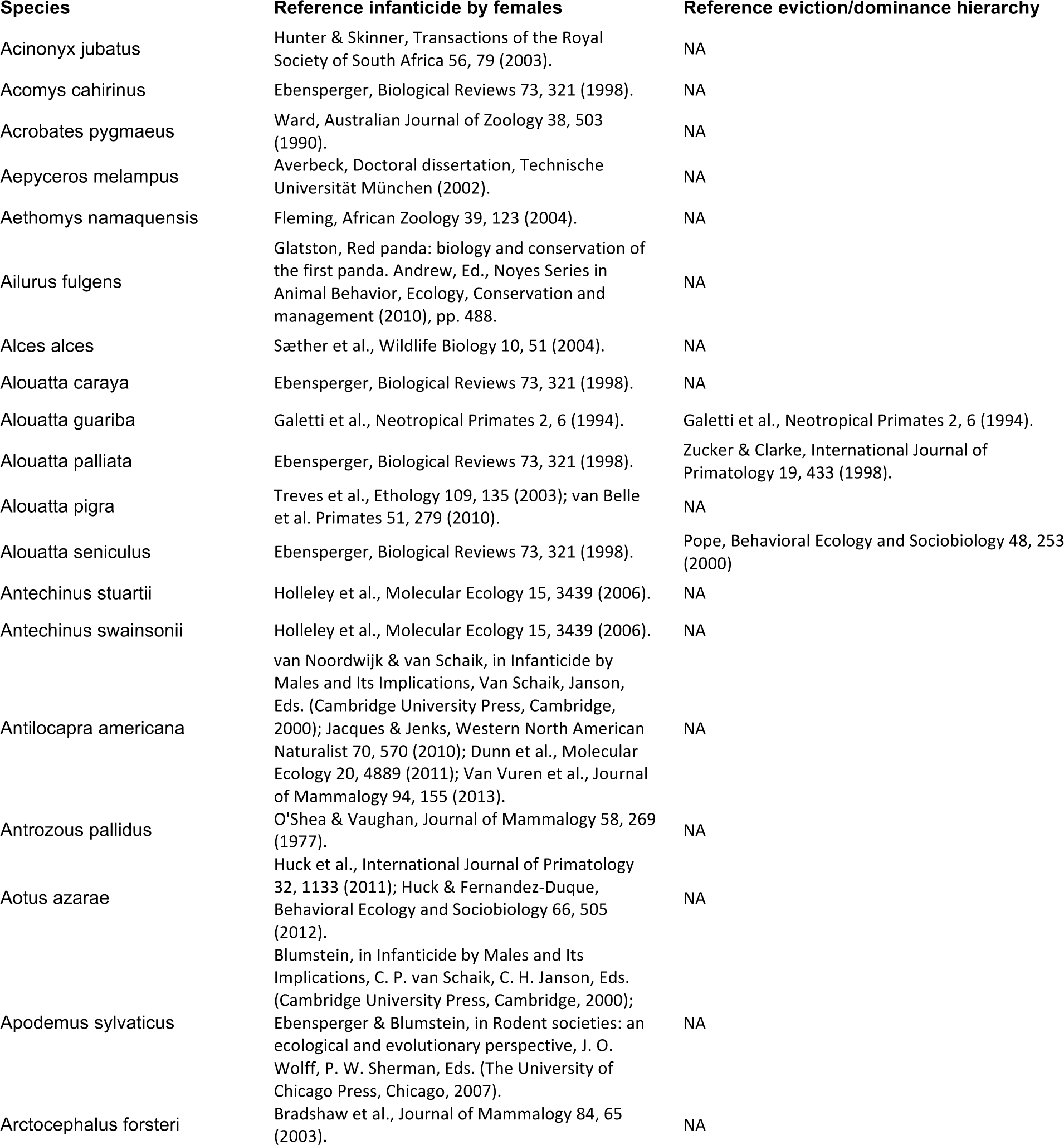

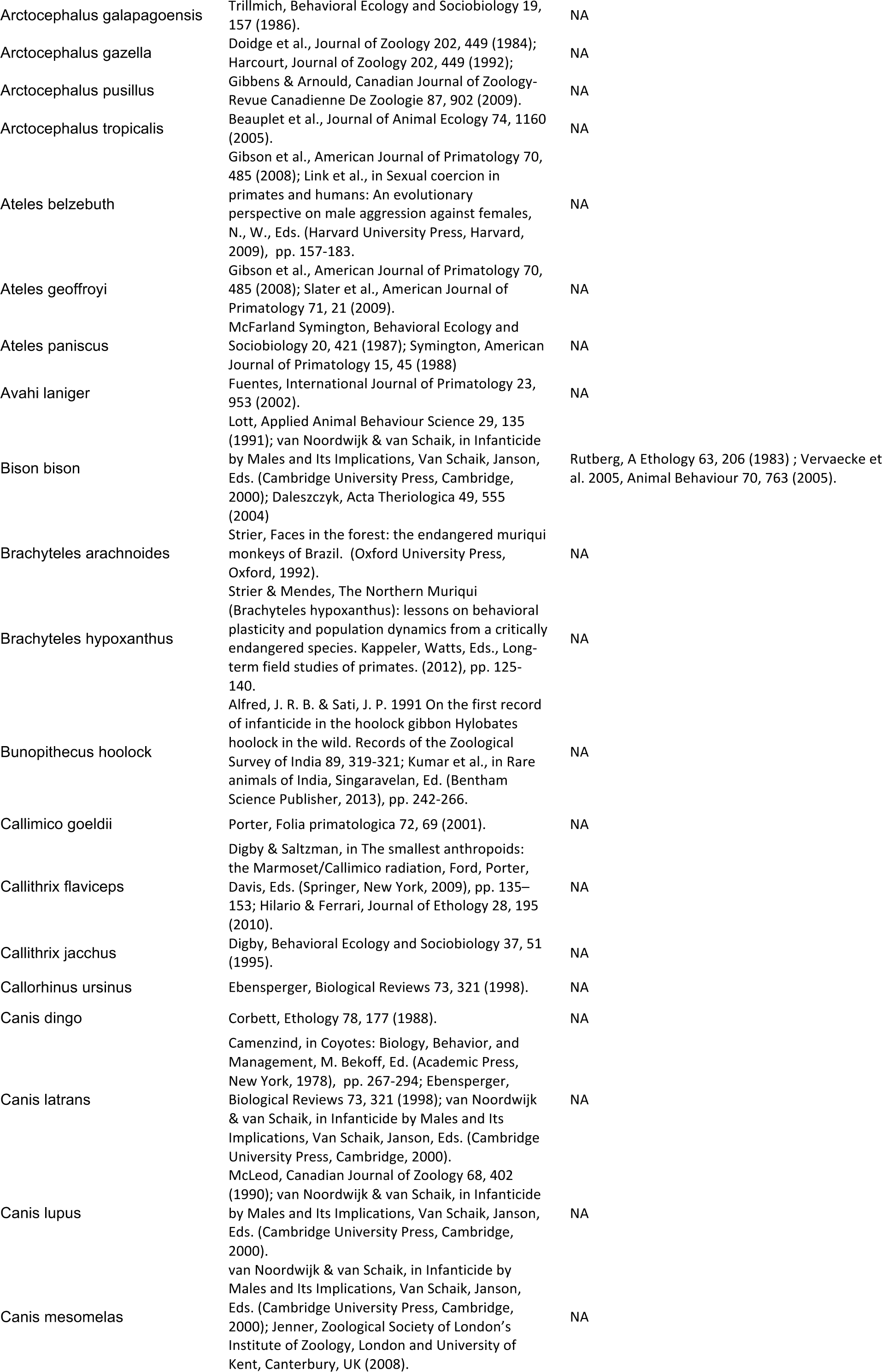

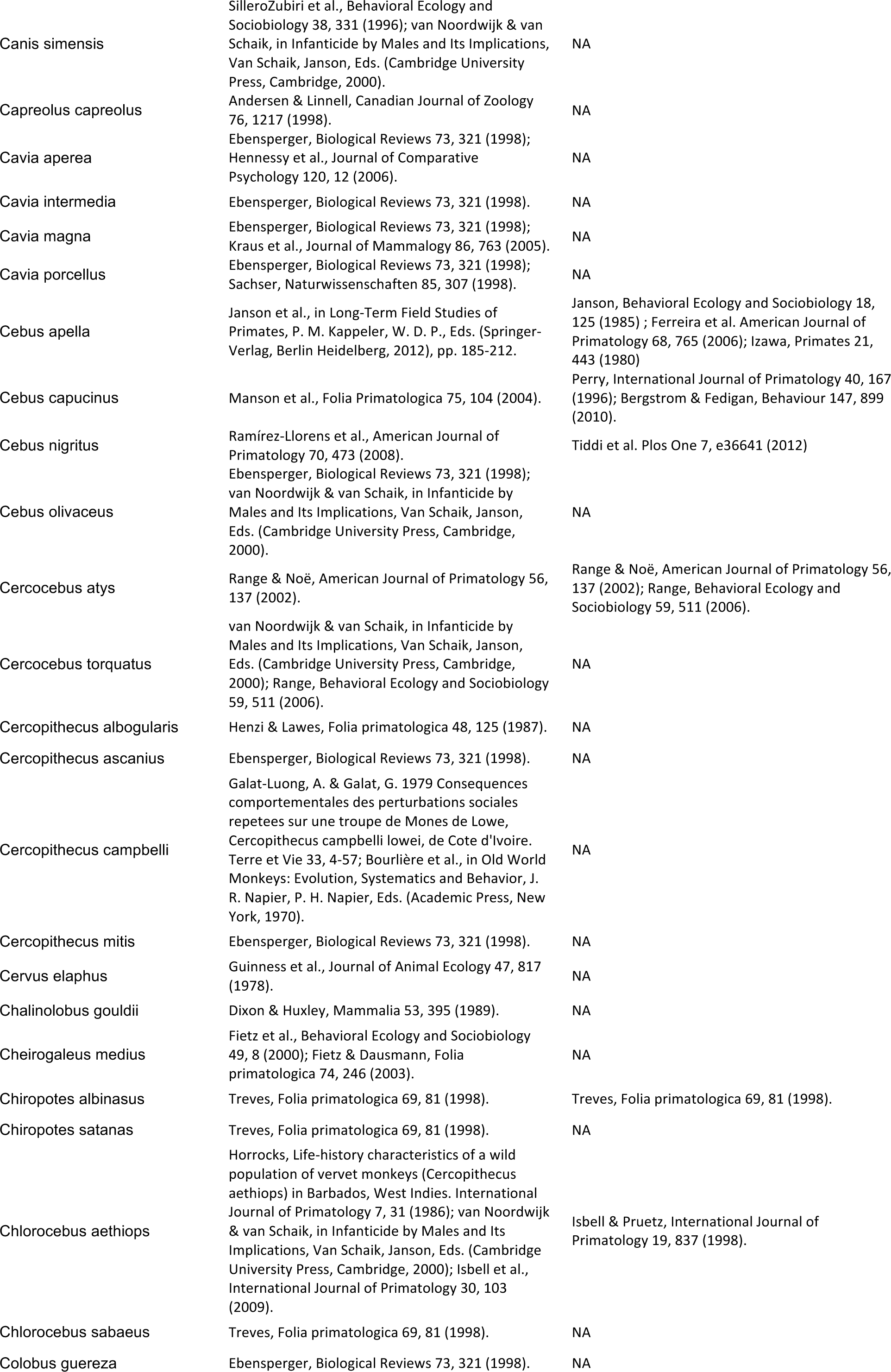

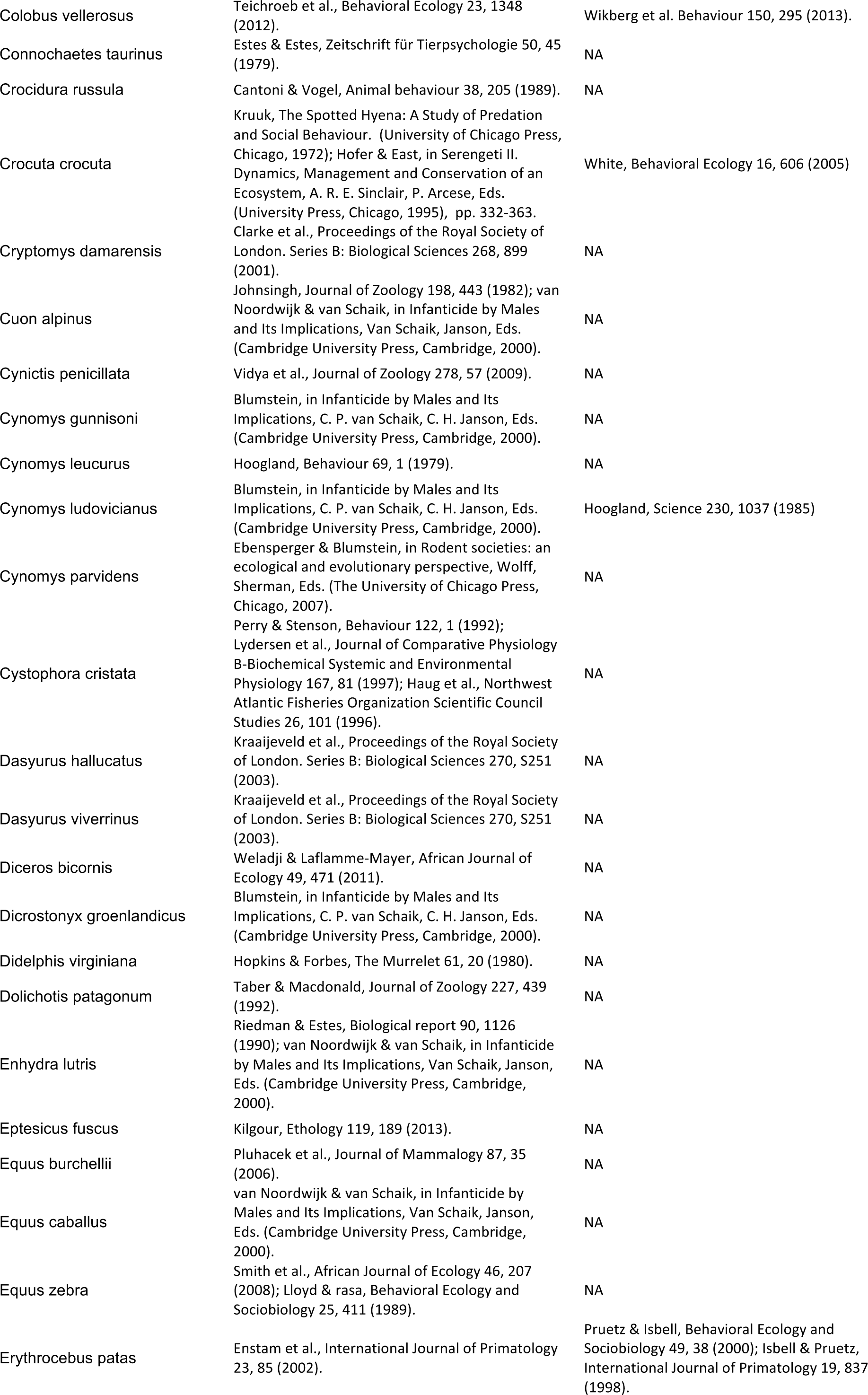

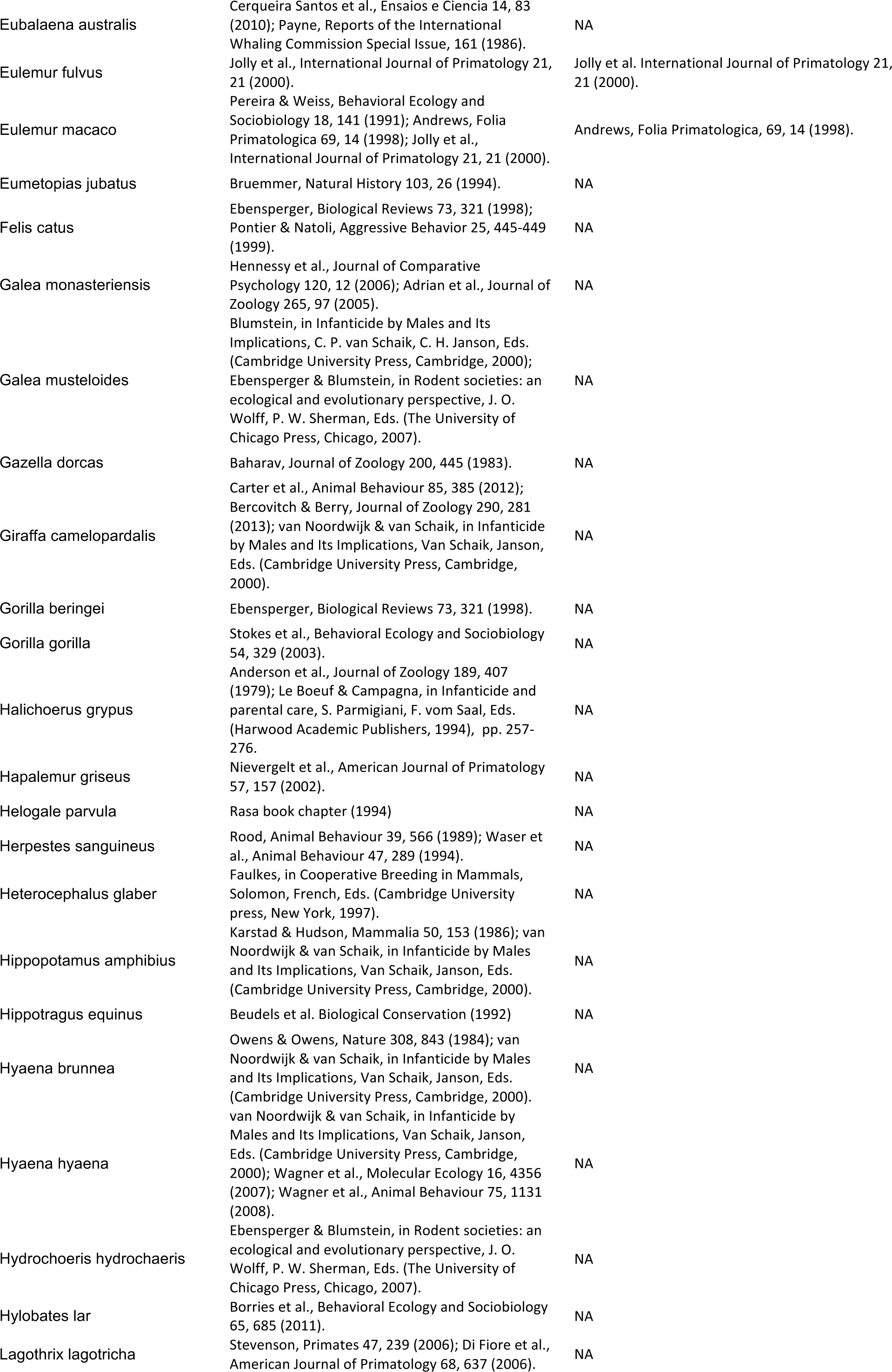

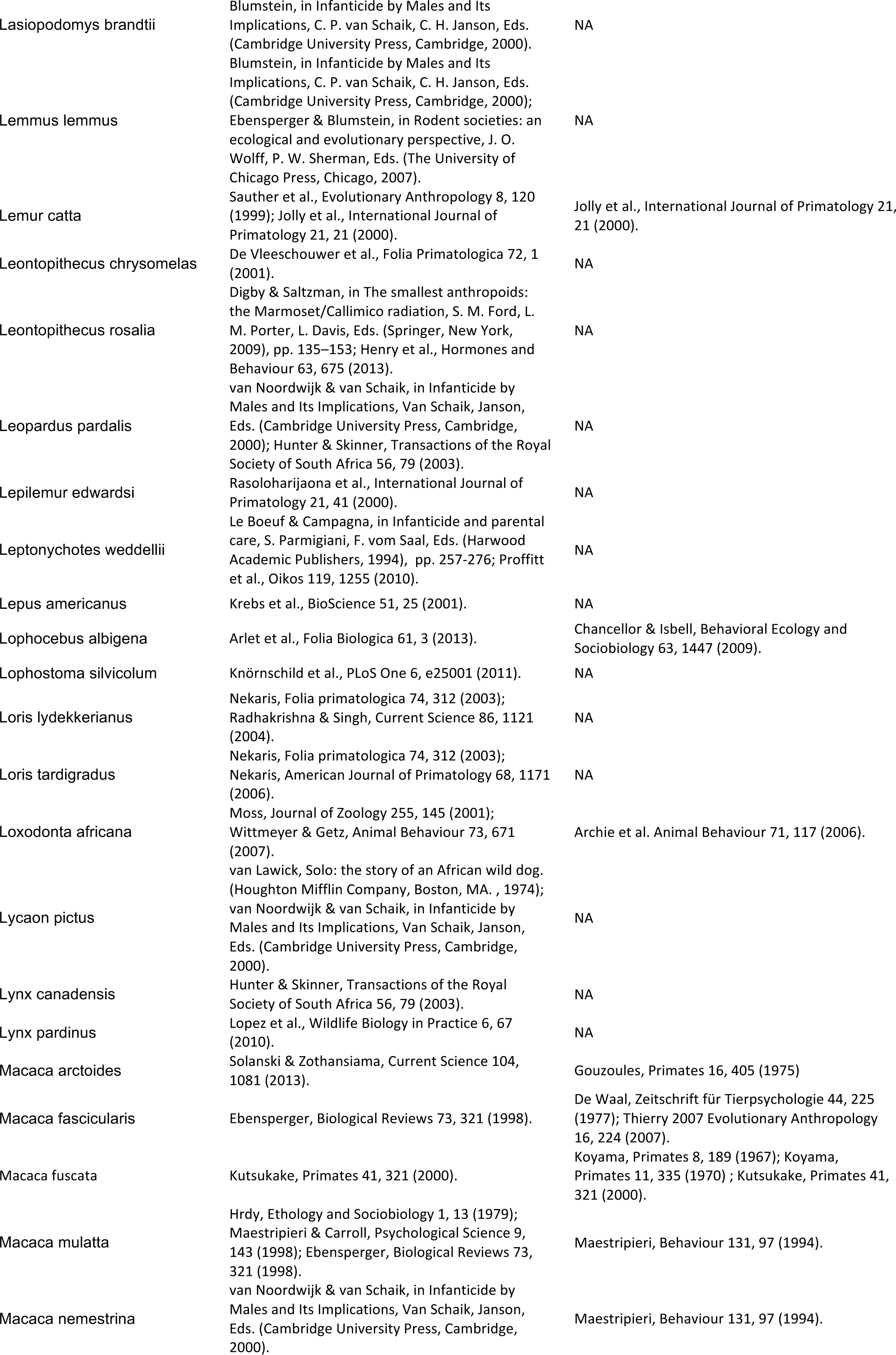

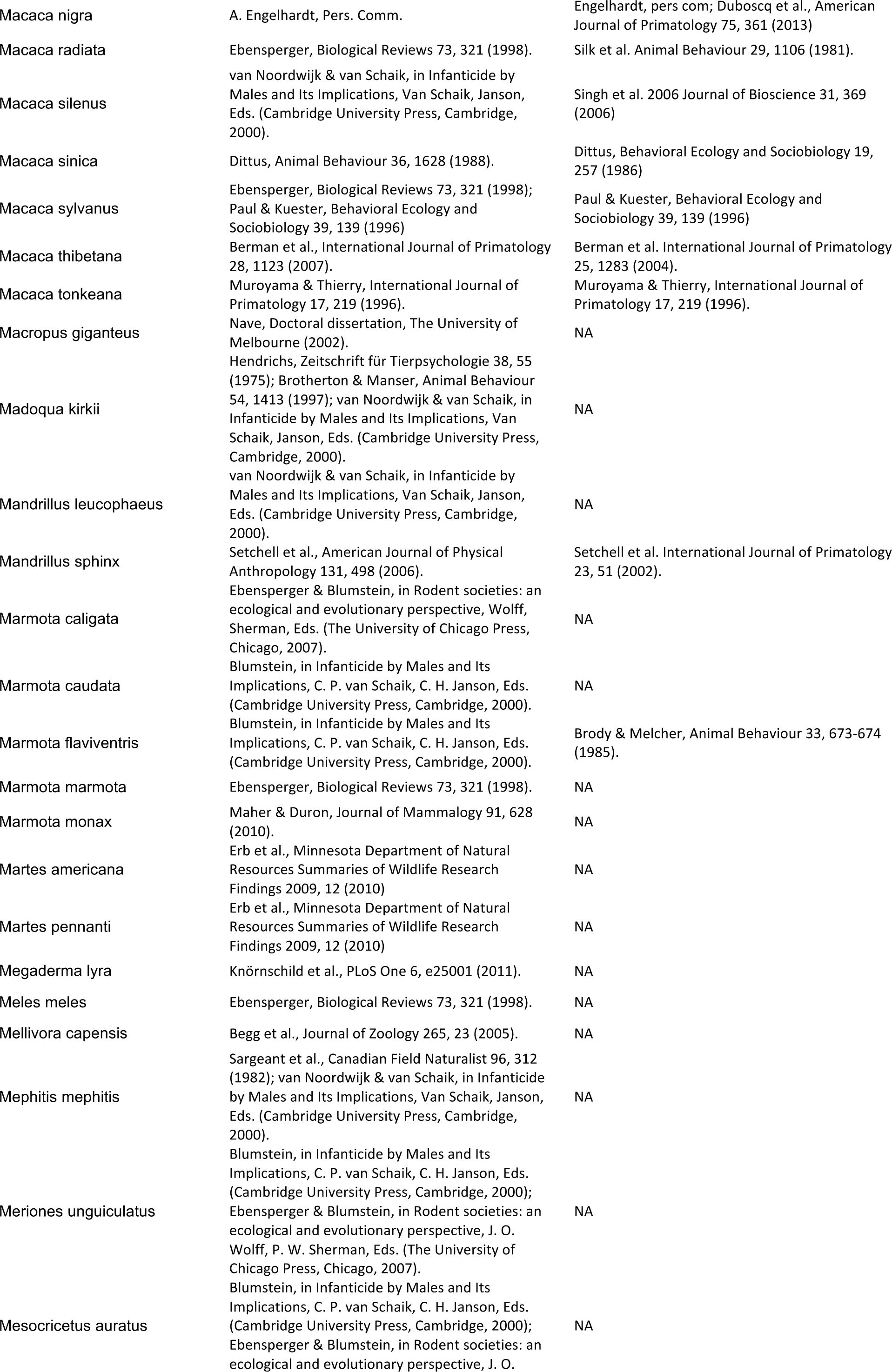

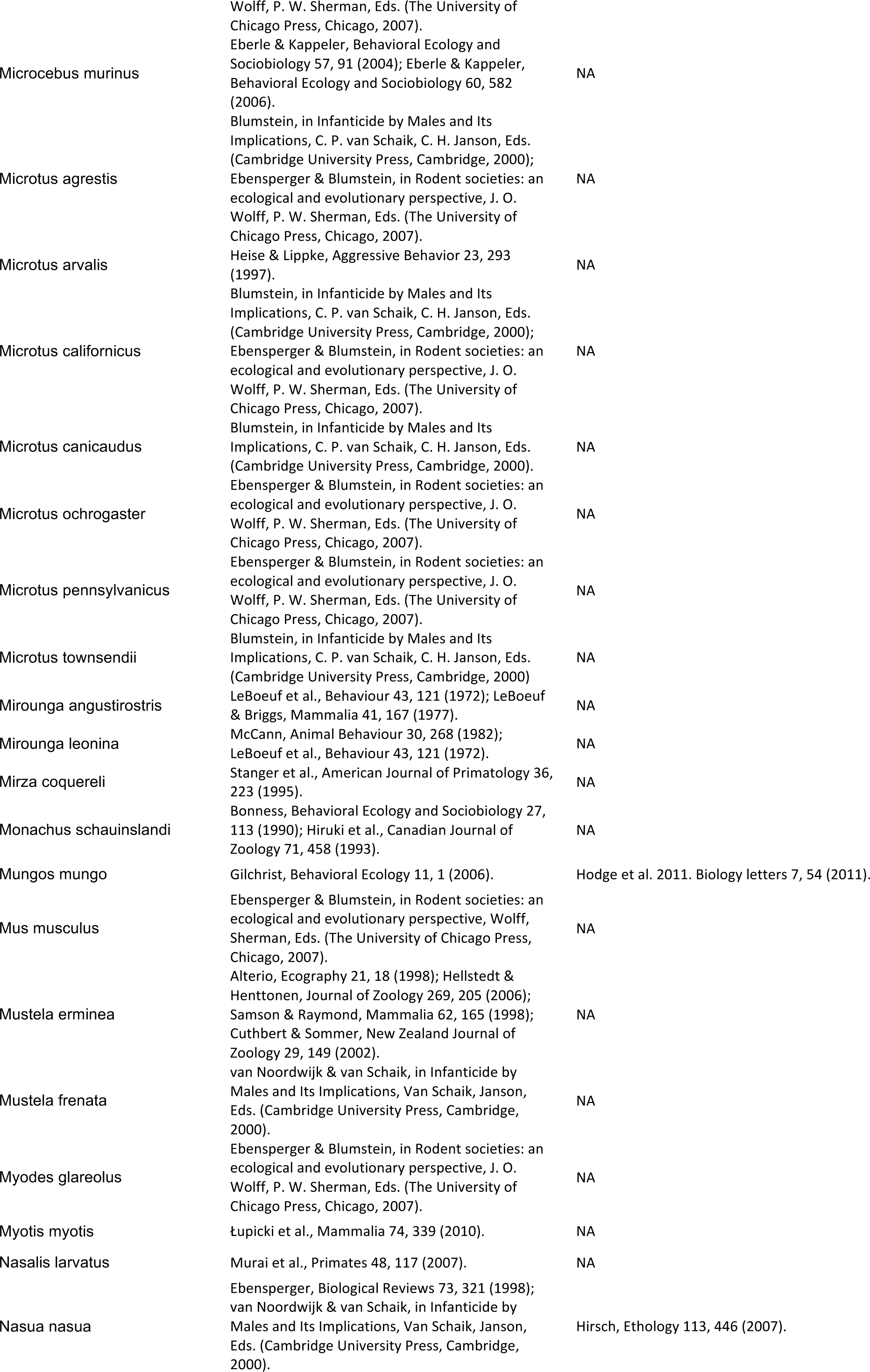

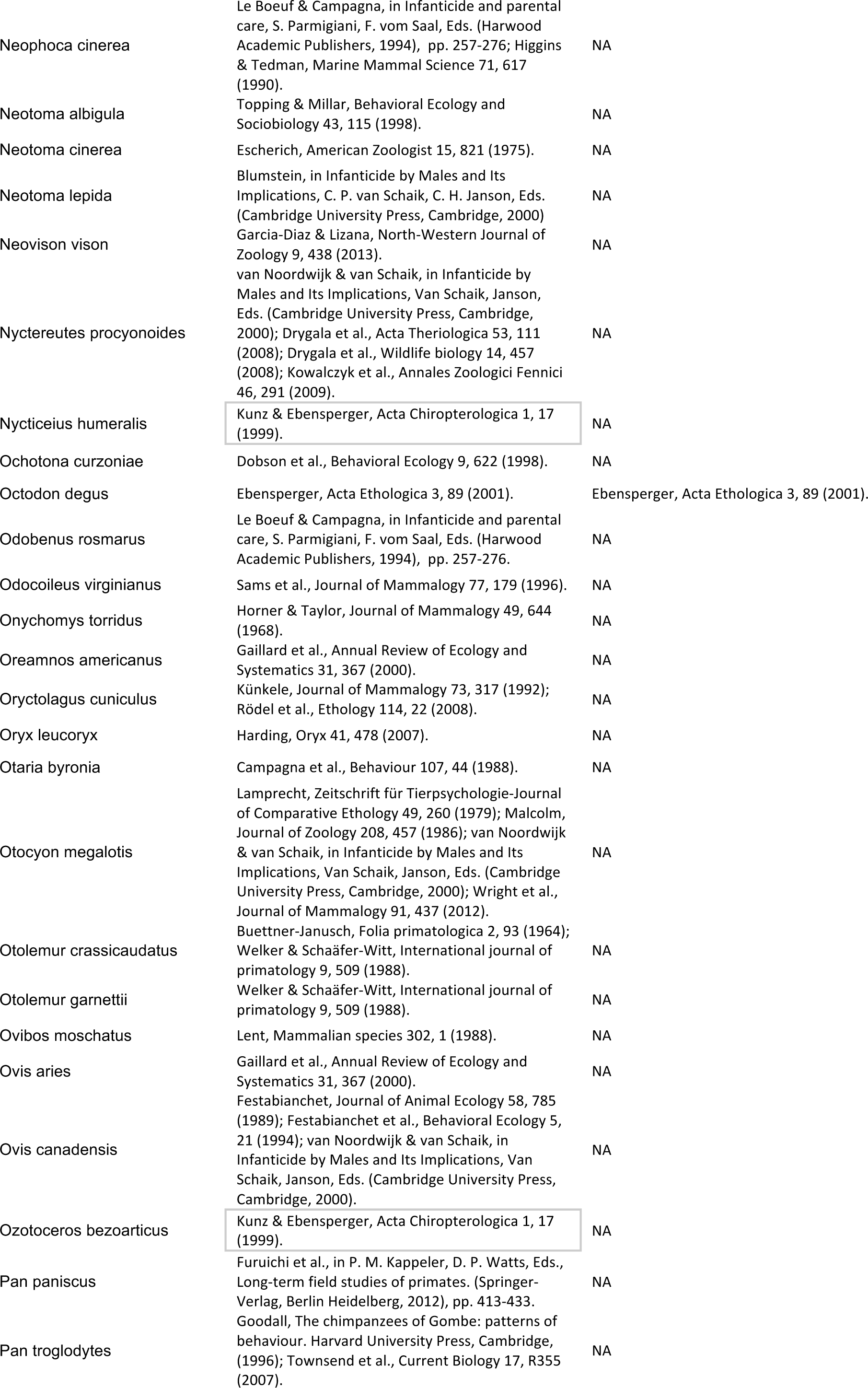

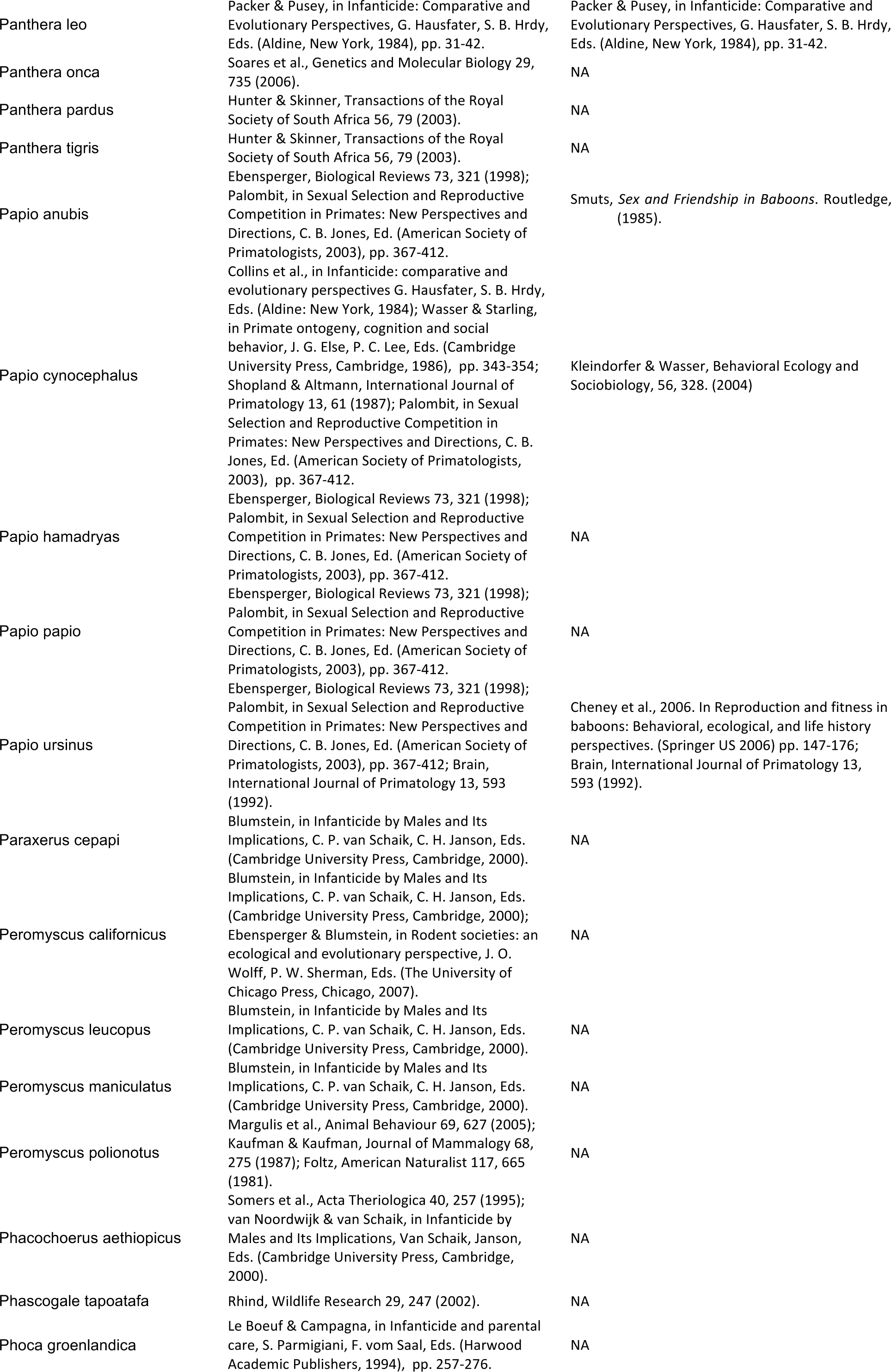

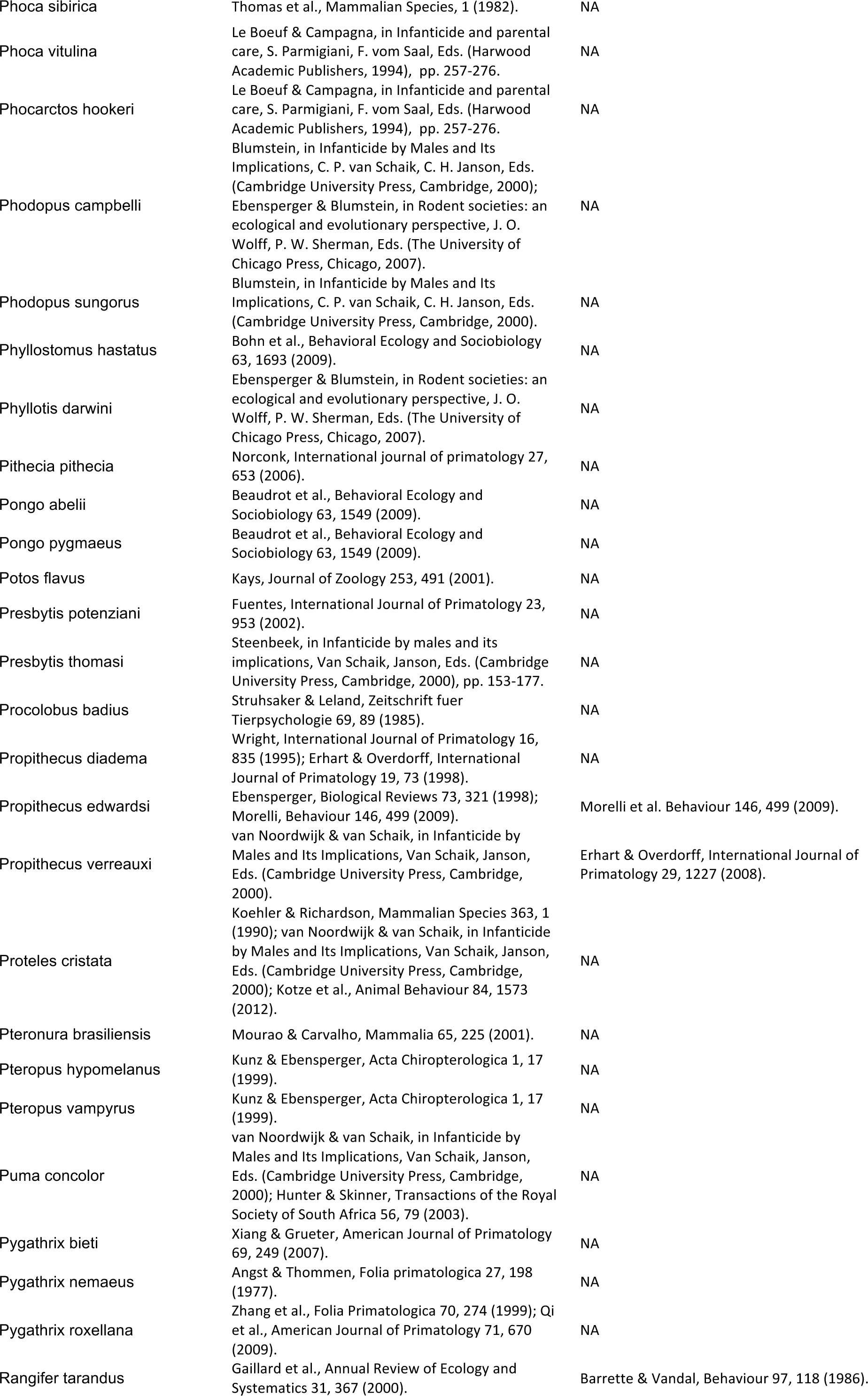

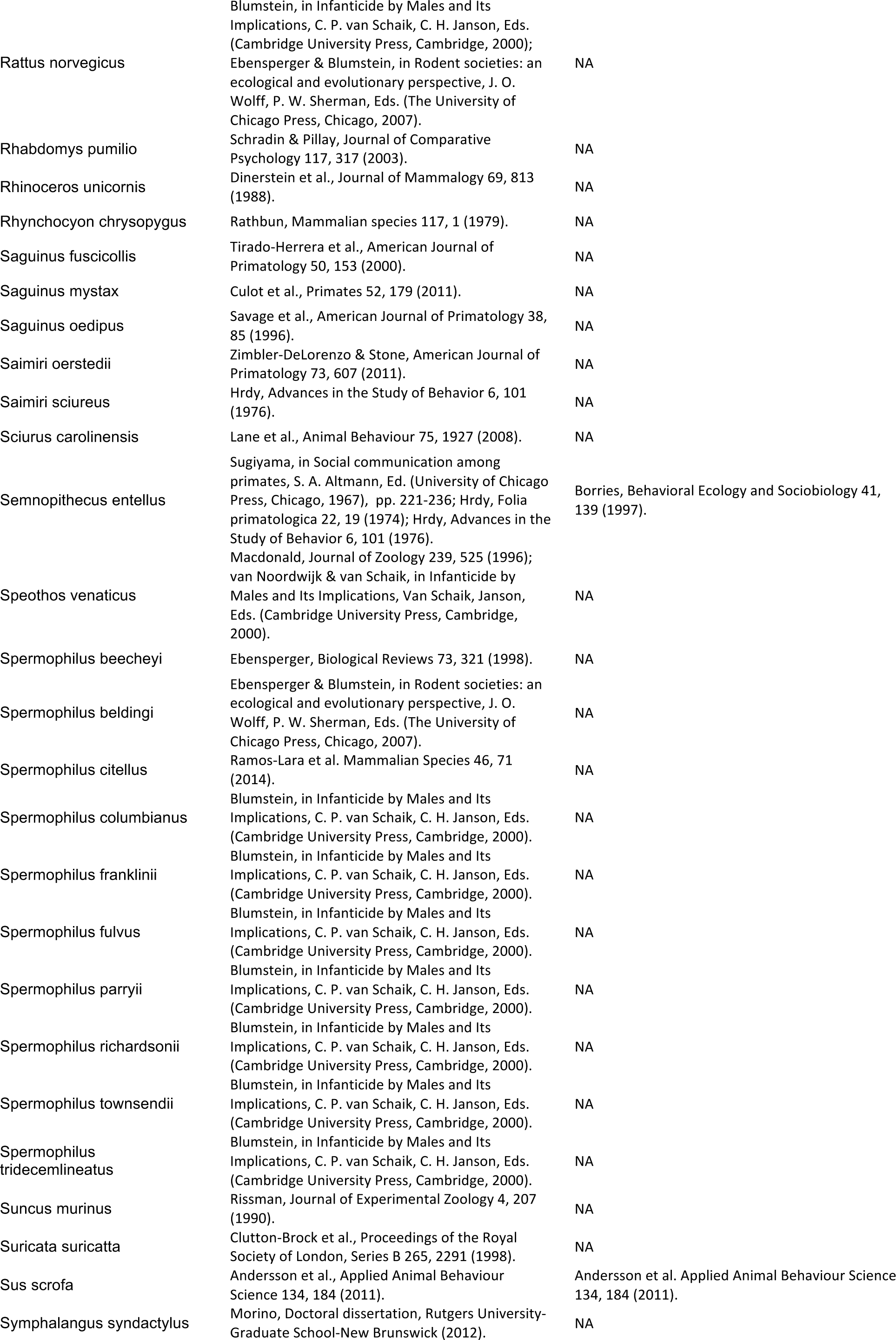

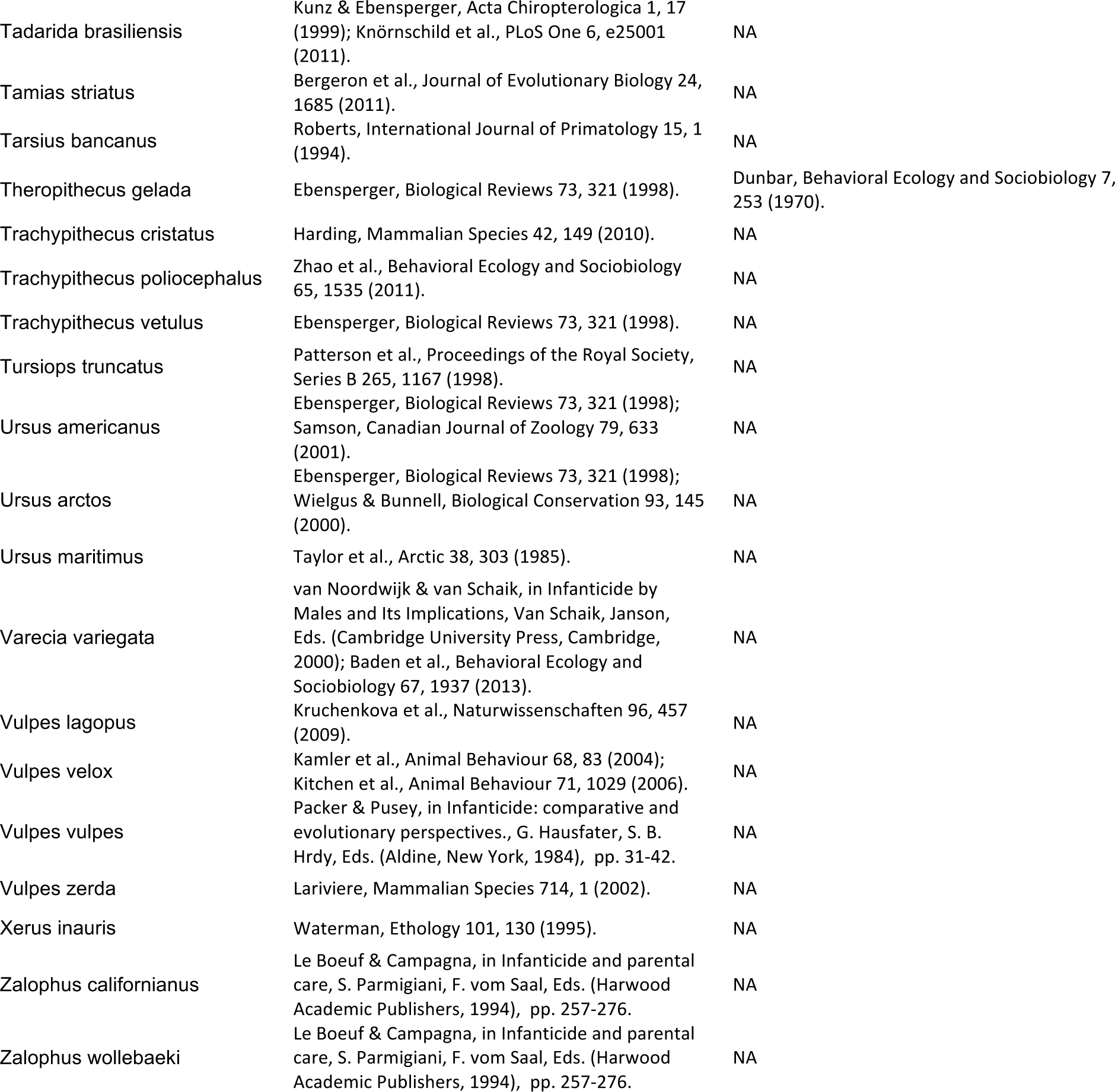

### B) Citations for population observation with details on female infanticide

The table lists the references with the observations on the characteristics of the killer and the victim and the circumstances of infanticide by females for the set of species for which this information was available. All references are listed in full below.

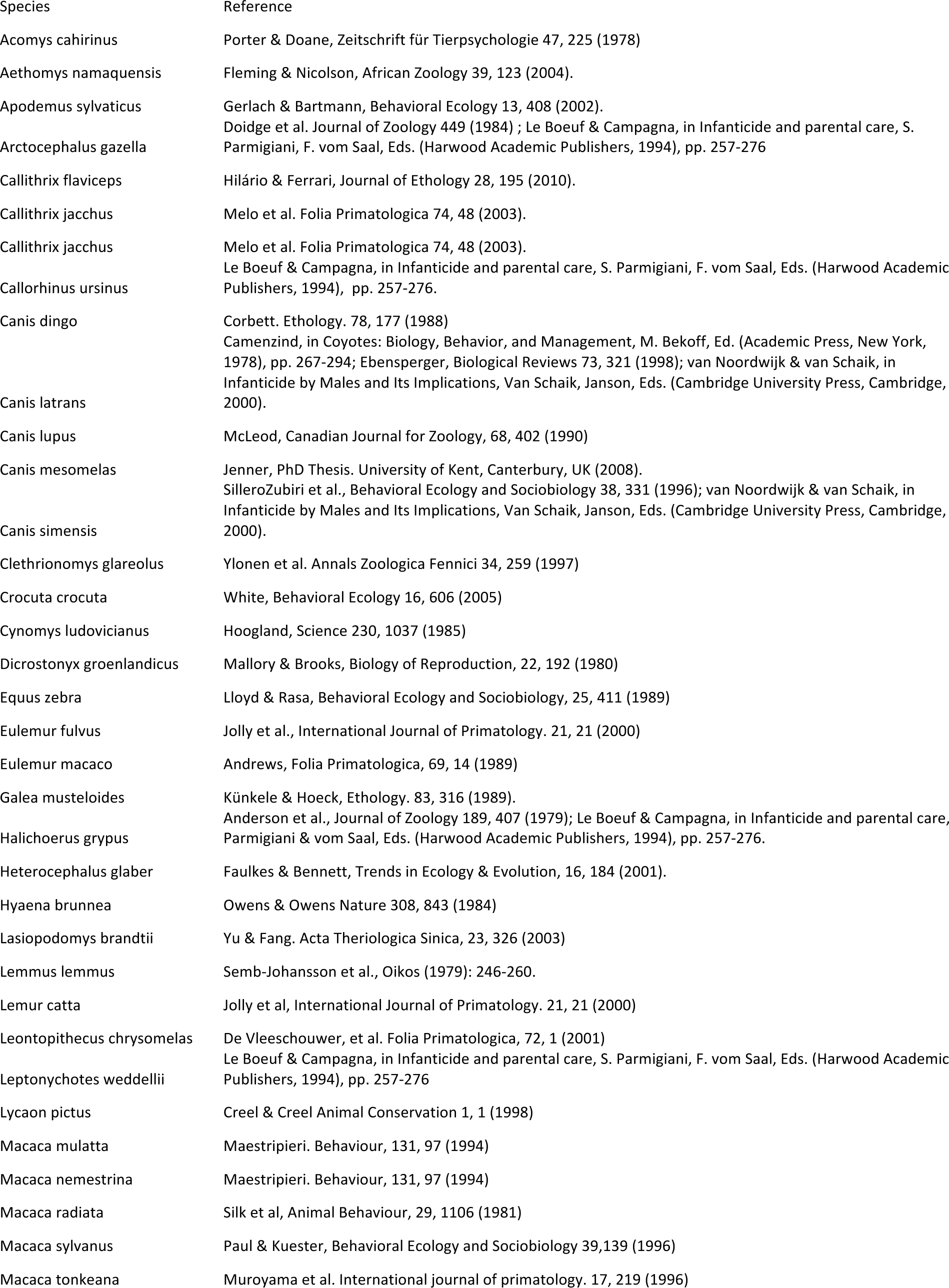

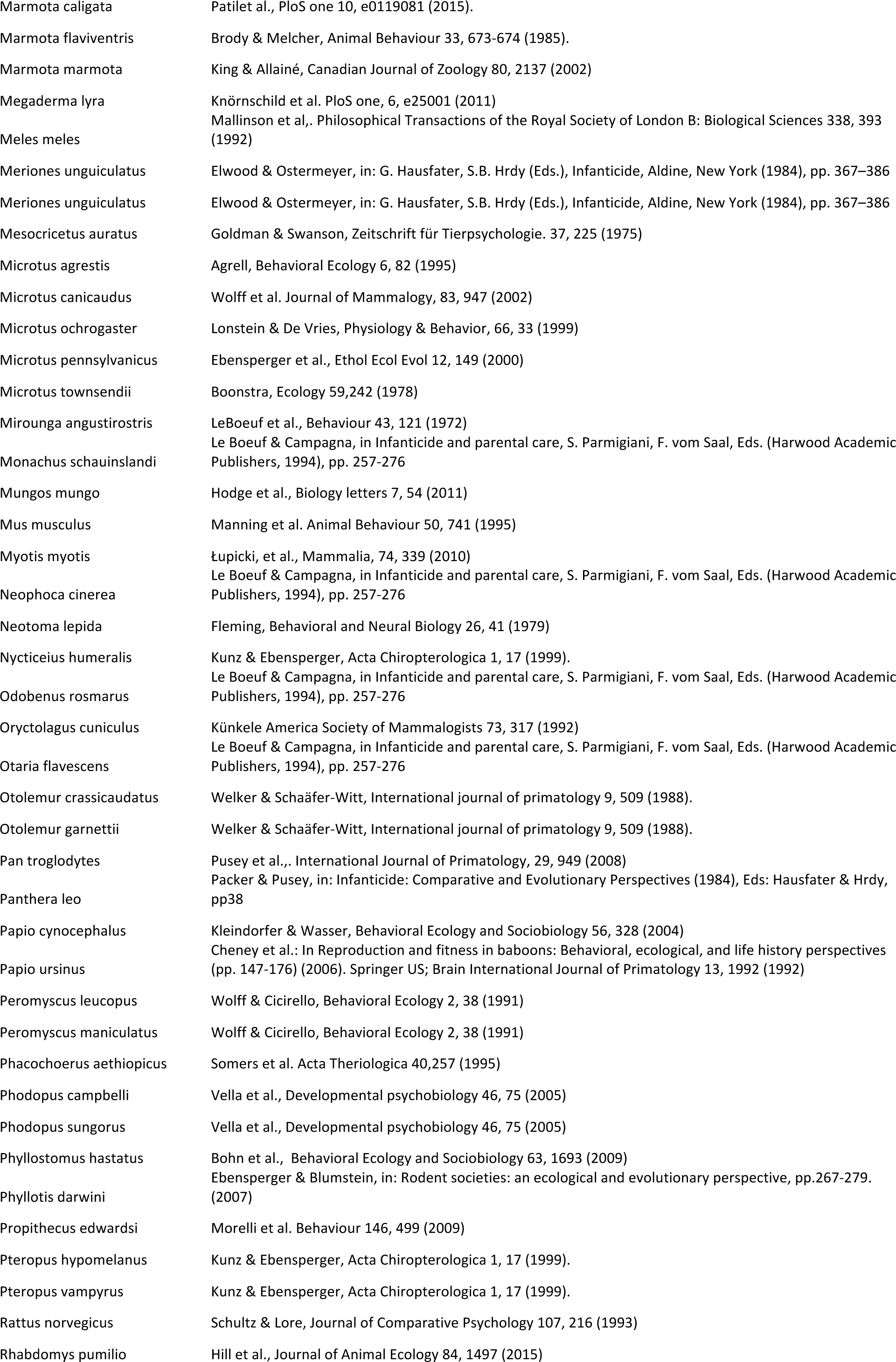

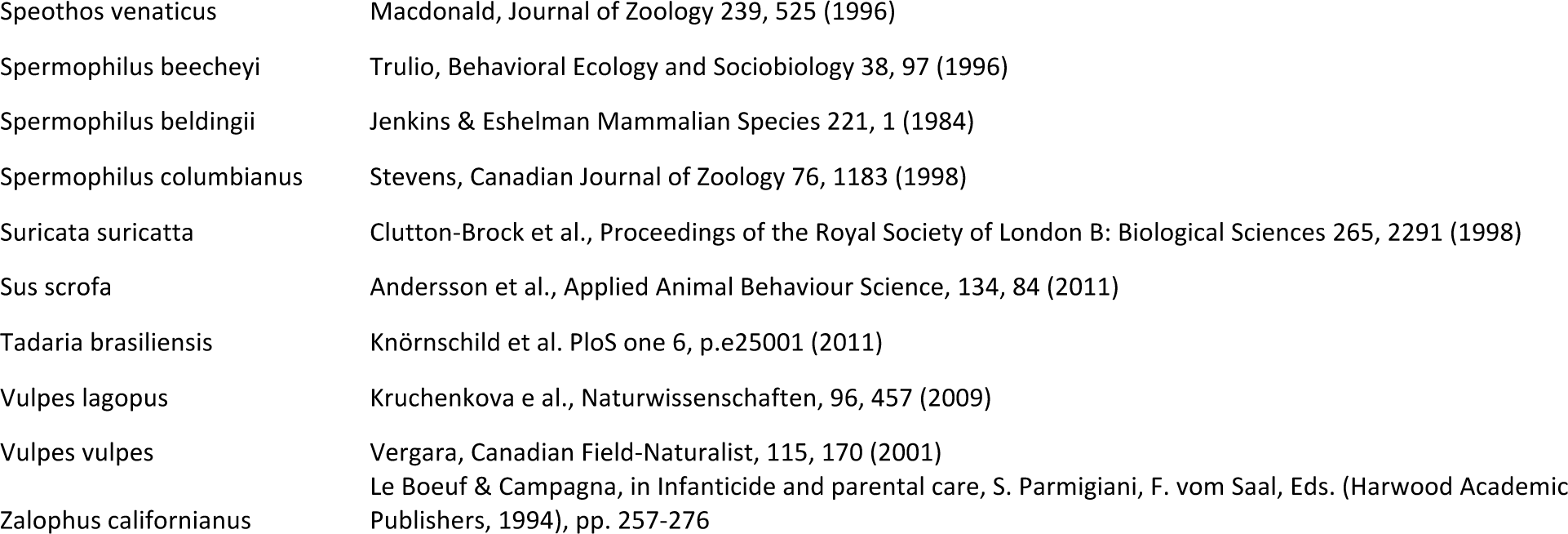

## Supplementary File 2 for Lukas & Huchard: The evolution of infanticide by females in mammals

Credit for animal drawings used in Figure 1

All drawings were downloaded from PhyloPic: http://phylopic.org

Picture information is listed as:

Common name listed in figure (taxon identifier for picture on PhyloPic): Author

### Starting from top, clockwise

Old World primate (*Papio*): Uncredited

Hare (*Leporidae*): Sarah Werning

Squirrel (*Sciuridae*): Catherine Yasuda

Marmot (*Marmota monax*): Michael Keesey

Mice (*Muridae*): Madeleine Price Ball

Marsupial (*Marsupialia*): Sarah Werning

Bat (*Chiroptera*): Michael Keesey

Mongoose (*Herpestoidae*): Michael Keesey

Felids (*Panthera*): Sarah Werning

Canids (*Canidae*): Michael Keesey

Bear (*Ursus*): Steven Traver

Seal & sealions (*Pinnipedia*): Steven Traver

Marten (*Meles*): Uncredited

Ungulate (*Cervus*): Steven Traver

Lemur (*Daubentonia*): Uncredited

Great ape (*Gorilla*): Michael Keesey

New World primate (*Cebus*): Sarah Werning

